# NewWave: a scalable R/Bioconductor package for the dimensionality reduction and batch effect removal of single-cell RNA-seq data

**DOI:** 10.1101/2021.08.02.453487

**Authors:** Federico Agostinis, Chiara Romualdi, Gabriele Sales, Davide Risso

## Abstract

**Summary:** We present *NewWave*, a scalable R/Bioconductor package for the dimensionality reduction and batch effect removal of single-cell RNA sequencing data. To achieve scalability, *NewWave* uses mini-batch optimization and can work with out-of-memory data, enabling users to analyze datasets with millions of cells.

**Availability and implementation:** *NewWave* is implemented as an open-source R package available through the Bioconductor project at https://bioconductor.org/packages/NewWave/

**Supplementary information:** Supplementary data are available at *Bioinformatics* online.

## 1 Introduction

Dimensionality reduction is a key step for the analysis of single-cell RNA-seq (scRNA-seq) data. Principal component analysis (PCA) is a simple and efficient method that can be employed for this step. However, it suffers from several drawbacks, e.g., it assumes that the data are Gaussian and does not allow to correct for technical variability and biases. While transforming the data (e.g., by running PCA on log-normalized counts) can ameliorate these problems, count-based factor analysis models often yield better low-dimensional data representations (Risso *et al*., 2018; Townes *et al*., 2019).

In particular, our recent method, ZINB-WaVE (Risso *et al*., 2018), uses a zero inflated negative binomial model to find biologically meaningful latent factors. Optionally, the model can remove batch effects and other confounding variables (e.g., sample quality), leading to a low-dimensional representation that focuses on biological differences among cells.

ZINB-WaVE has been shown to be among the top performing methods in recent benchmarks (Sun *et al*., 2019; Raimundo *et al*., 2020). However, its main drawback is the lack of scalability, due to large memory requirements that prevent its use with more than a few cores. To address this, we have re-implemented the model of ZINB-WaVE in a new Bioconductor package, *NewWave*, which allows users to massively parallelize computations using PSOCK clusters. Here, we show that *NewWave* is able to achieve the same, or even better, performance of ZINB-WaVE at a fraction of the computational speed and memory usage, reducing the runtime by 90% with respect to ZINB-WaVE.

## 2 Software implementation

*NewWave* uses a factor analysis framework similar to that of ZINB-WaVE (Risso *et al*., 2018), with the important difference that the gene-level read counts are assumed to come from a negative binomial distribution without zero inflation. In fact, the majority of large scRNA-seq data use unique molecular identifiers (UMIs) and UMI data are not zero inflated (Townes *et al*., 2019; Svensson, 2020). Briefly, the log of the expected value of the read count matrix is modeled as a regression of three terms: known cell covariates (*X*, e.g., batch), known gene covariates (*V*, e.g., an intercept with the role of normalization) and latent factors (*W*) that define a low-dimensional space that describe the unknown biological signal (Fig. 1A and Supplementary Information). With a high number of cells, these matrices are large and it may not be easy to control how many times they are copied during parallel execution.

**Fig. 1.**
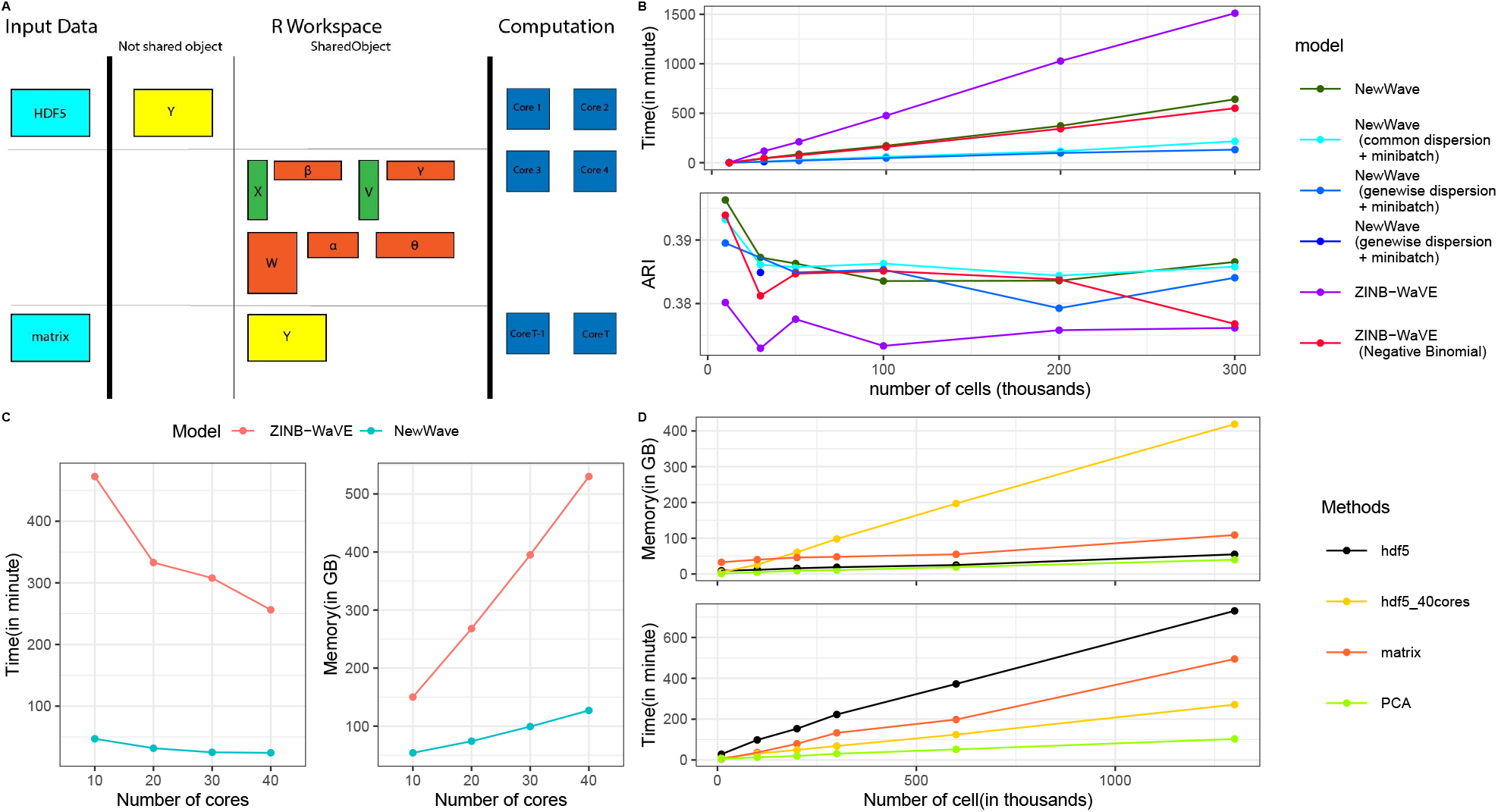
Implementation and performance of NewWave. Unless otherwise noted, we used 10% of the observations as the size of the mini-batches and 10 cores. A. Schema of the NewWave model, indicating which matrices are in shared memory (see Supplementary Information for more details). B. Speed (top) and Adjusted Rand Index (ARI, bottom) of NewWave (in-memory data) with different choices of the parameters and ZINB-WaVE applied to the BICCN dataset (Yao et al., 2020) with a maximum of 312,000 cells and after selecting the 1,000 most variable genes. The reported ARI is computed as the mean ARI of 100 k-means clustering procedures with the number of centroids set to the known number of labels (*k* = 20). C. Speed and RAM usage of NewWave using a subset of 100,000 cells varying the number of cores used for computation. D. RAM usage (top) and speed (bottom) of NewWave on the 10X 1.3M cell datasets with 1,000 most varable genes.

The three main strategies that *NewWave* uses to limit the computational problems of working with large matrices are: (i) the use of shared memory objects in PSOCK clusters to avoid redundant data copies, (ii) the use of mini-batch optimization algorithms to speed-up computations, and (iii) the use of out-of-memory data representations (such as HDF5 files) to limit memory usage.

The optimization procedure can be represented as a cycle of three steps, iterated until convergence: (i) optimization of the dispersion parameters (either common dispersion or gene-wise dispersion); (ii) optimization of gene-wise parameters; (iii) optimization of cell-wise parameters.

One of the main advantages of our model specification is that it naturally results in an embarrassingly parallel task. In fact, except for the optimization of the global dispersion parameter (common to all genes), all the steps use only one gene (cell) at a time for the optimization of gene (cell) parameters. In addition to parallelization, this setup is ideal for mini-batch optimization strategies. At any one step, we can use a random subset of cells (genes) to estimate the gene (cell) parameters.

On-disk datasets are managed through the *DelayedArray* package (Pagès *et al*., 2019), which allows block processing and delayed operations on data stored in HDF5 files. While all covariates and parameter matrices are stored in shared memory among child processes, the input data can reside either in shared memory or on-disk as an HDF5 file (Fig. 1A).

## 3 Results and discussion

The application of NewWave to subsamples of large datasets, in particular when relying on mini-batches, shows a better scalability than ZINB-WaVE without loss of accuracy (Fig. 1B; see Supplementary Information for details on the analysis). Strikingly, the negative binomial model outperforms its zero-inflated counterpart, confirming that this is a preferable model for UMI data (Townes *et al*., 2019; Svensson, 2020).

In addition to speed, we measured the scalability of *NewWave* in terms of RAM usage (Fig. 1C, D). As expected, there is a speed-RAM trade-off when using data in-memory or on-disk. Runtimes increase when using HDF5, due to the additional I/O, but this dramatically decreases the RAM consumption (Fig. 1D). This in turns allows the use of more cores. Using 40 cores, the computational time of our HDF5 implementation is lower than that of of the in-memory data with 10 cores, allowing us to analyze of 1.3M cells in 271 minutes using 109GB of RAM.

*NewWave* is available as an open-source package through the Bioconductor project. The package includes a vignette with a tutorial. In addition, the code to reproduce all the analyses presented here is available at https://github.com/fedeago/NewWave-script.

## Supplementary material

### 1 Model specification

NewWave assumes a negative binomial distribution for the expression data. Given *n* samples and *J* genes across all samples and denoting by *Y*_*i j*_ the count of gene *j* in cell *i*, the likelihood function for the observed count *y*_*i j*_ is

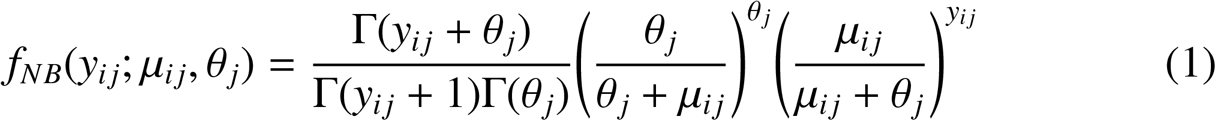

in this parametrization the variance is

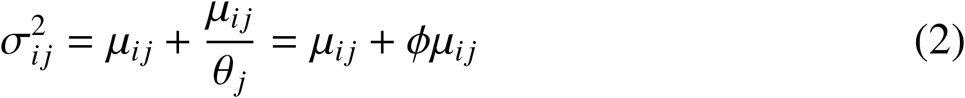

We specify the following regression model:

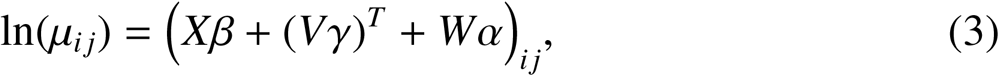

where *X* is a matrix with dimension *n* × *M* where *M* is the number of cell-level covariates. It typically contains a set of dummy variables that specify the batch of the samples and by default includes an intercept. *V* is a matrix with dimension *J* × *L* where *L* is the number of gene-level covariates. It typically only contains an intercept, used as library size normalization. *W* is a matrix with dimension *n* × *K*, where *K* is the number of latent factors, and it contains the low-rank representation of the cells. The parameters *α, β*, and *γ* are the regression coefficients for *W, X*, and *W*, respectively.

We note that (3) is a special case of the model of Risso et al. (2018) and we refer the reader to that paper for additional details.

### 2 Parameter estimation

The log-likelihood function to be maximized is

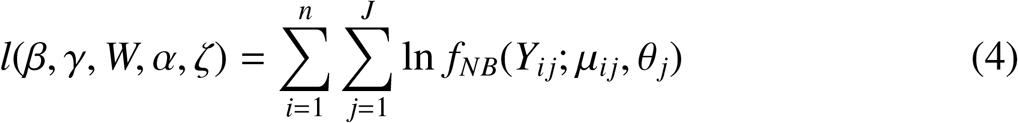

The estimation of the parameters is done with a penalized approach for *β, α, W, γ* and not penalized for *ζ*:

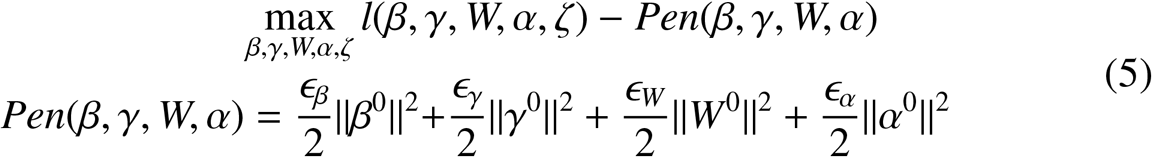

this penalization prevents overfitting (Risso et al., 2018).

Our iterative estimation procedure follows closely from Risso et al. (2018), considering the special case of no zero inflation.

### 2.1 Dispersion parameters

Special attention must be paid to the estimation of the dispersion parameters *θ* _*j*_. These can be estimated in a gene-wise fashion, or assuming a common dispersion parameter *θ* _*j*_ = *θ* for all the genes.

This choice yields two different implementations. When the user wants to estimate a common dispersion parameter, our implementation takes advantage of the mini-batch strategy outlined below. However, the computation cannot leverage parallel computations, since only one value needs to be estimated. On the other hand, when the user requires gene-wise dispersion parameters, the computations can proceed in parallel for each gene, but the mini-batch strategy is not effective, as it slows down computations.

### 2.2 Mini-batches

Even when the mini batch approach is chosen, the first iteration will be done using all the observation; this method shows better performance in terms of time needed to converge. If the common dispersion approach is chosen, starting from the second iteration, only a random subset *b* of size *m* of the observations is used at each iteration to estimate the parameter. Note that the parameter estimate is assumed to describe the variation across all observations, even those that do not belong to *b*.

At each iteration, the estimated parameter is optimal for *b*, but there is no guarantee of its optimality on the full data. Hence, the value of the parameter estimate is updated only if it leads to an increase in the log-likelihood function in the full data.

The optimization of the cell- and gene-specific parameters is performed with a mini-batch approach after a first iteration that uses all data. When parallel computing is used, each child process uses only *m/c* observations where *m* is the number of observations in the mini-batch and *c* is the number of child processes.

### 3 Details on the benchmark

#### 3.1 Compared strategies

We benchmarked different estimation strategies implemented in the *Newwave* package and we compared their results with those of ZINB-WaVE, both with and without zero inflation.

The estimation strategies in *Newwave* are

- **Default (Common dispersion, no mini-batches)**. A single dispersion parameter, common to all genes, is estimated. All cells and genes are used for the estimation of gene- and cell-specific parameters.
- **Common dispersion** + **mini-batch**. A single dispersion parameter, common to all genes, is estimated. Mini-batches of 10% total cells and *m* = 100 genes are used to estimated for the estimation of gene- and cell-specific parameters, respectively.
- **Gene-wise dispersion** + **mini-batch**. Gene-wise dispersion parameters are estimated. Mini-batches of 10% total cells and *m* = 100 genes are used to estimated for the estimation of gene- and cell-specific parameters, respectively.

#### 3.2 Datasets used

We used two publicly available datasets for our benchmark, namely the BICCN dataset and the 10X Brain dataset.

##### BICCN data

We created the first dataset starting from the mouse primary motor cortex datasets generated by the BRAIN Initiative Cell Census Network (BICCN) (Yao et al., 2020), which we refer to as the BICCN dataset.

The raw data was generated as part of the BICCN consortium and can be downloaded from http://data.nemoarchive.org/biccn/lab/zeng/transcriptome/.

The known batch effect present in this dataset is due to the difference between platforms, namely 10X Chromium single-cell sequencing v2 and v3 and 10X Chromium single-nucleus sequencing.

We selected the 1,000 most variable genes after correcting for the batch effects using the mutual nearest neighbor method (Haghverdi et al., 2018), as implemented in the *fastMNN* function of the *batchelor* Bioconductor package (v. 1.3.14). All the methods were applied to these 1,000 genes.

##### 10X Brain data

The second dataset is the 1.3 million brain cell single-cell RNA-seq (scRNA-seq) data set generated by 10X Genomics without known batch effect.

The dataset is available in dense HDF5 format as part of the *TENxBrainData* Bioconductor package at https://bioconductor.org/packages/TENxBrainData. We selected the 1,000 most variable genes on the entire dataset.

## 4 Acknowledgements

The authors thank Hongkui Zeng and the members of the BICCN consortium for sharing the BICCN dataset.

## 5 Supplementary Tables

**Table 1:** Time, Akaike Information Criterion (AIC), and Adjusted Rand Index (ARI) for the different dimentionality reduction methods on the BICCN dataset. This table reports the data displayed in Fig. 1B.

| Model | # of cells | Time in minutes | AIC | ARI |
| --- | --- | --- | --- | --- |
| NewWave | 10000 | 15.64 | 39097804 | 0.3963 |
| NewWave (common dispersion + minibatch) | 10000 | 5.67 | 39177295 | 0.3932 |
| NewWave (genewise dispersion + minibatch) | 10000 | 3.96 | 38933804 | 0.3895 |
| ZINB-WaVE | 10000 | 44.06 | 43068497 | 0.3802 |
| ZINB-WaVE (Negative Binomial) | 10000 | 16.24 | 50010784 | 0.3939 |
| NewWave | 30000 | 2715.70 | 116840934 | 0.3873 |
| NewWave (common dispersion + minibatch) | 30000 | 618.29 | 117173555 | 0.3861 |
| NewWave (genewise dispersion + minibatch) | 30000 | 679.45 | 116363662 | 0.3849 |
| ZINB-WaVE | 30000 | 7036.46 | 128899425 | 0.3730 |
| ZINB-WaVE (Negative Binomial) | 30000 | 2659.80 | 149574224 | 0.3812 |
| NewWave | 50000 | 5155.36 | 194632320 | 0.3863 |
| NewWave (common dispersion + minibatch) | 50000 | 1668.14 | 195018975 | 0.3857 |
| NewWave (genewise dispersion + minibatch) | 50000 | 1274.62 | 193863319 | 0.3849 |
| ZINB-WaVE | 50000 | 12697.25 | 214638707 | 0.3775 |
| ZINB-WaVE (Negative Binomial) | 50000 | 4487.82 | 249048748 | 0.3847 |
| NewWave | 100000 | 10288.96 | 389698453 | 0.3835 |
| NewWave (common dispersion + minibatch) | 100000 | 3617.65 | 390483587 | 0.3863 |
| NewWave (genewise dispersion + minibatch) | 100000 | 2827.11 | 388019619 | 0.3853 |
| ZINB-WaVE | 100000 | 28576.99 | 430023667 | 0.3733 |
| ZINB-WaVE (Negative Binomial) | 100000 | 9515.64 | 498933752 | 0.3851 |
| NewWave | 200000 | 22358.31 | 780334158 | 0.3836 |
| NewWave (common dispersion + minibatch) | 200000 | 6970.93 | 782011394 | 0.3844 |
| NewWave (genewise dispersion + minibatch) | 200000 | 5923.64 | 777041944 | 0.3793 |
| ZINB-WaVE | 200000 | 61668.90 | 861598414 | 0.3758 |
| ZINB-WaVE (Negative Binomial) | 200000 | 20600.85 | 999580927 | 0.3838 |
| NewWave | 300000 | 38478.19 | 1211144264 | 0.3866 |
| NewWave (common dispersion + minibatch) | 300000 | 12992.71 | 1213490638 | 0.3858 |
| NewWave (genewise dispersion + minibatch) | 300000 | 7919.25 | 1205982980 | 0.3841 |
| ZINB-WaVE | 300000 | 90670.66 | 1338439179 | 0.3762 |
| ZINB-WaVE (Negative Binomial) | 300000 | 33036.59 | 1551562402 | 0.3768 |

**Table 2:** Time and RAM usage of NewWave and ZINB-WaVE varying the number of CPUs. This table reports the data displayed in Fig. 1C.

| **Model** | **Time in minutes** | **#of cores** | **RAM in in GB** |
| --- | --- | --- | --- |
| NewWave | 46.97 | 10 | 54 |
| NewWave | 31.69 | 20 | 74 |
| NewWave | 25.00 | 30 | 99 |
| NewWave | 24.22 | 40 | 127 |
| ZINB-WaVE | 472.53 | 10 | 150 |
| ZINB-WaVE | 332.94 | 20 | 268 |
| ZINB-WaVE | 307.83 | 30 | 395 |
| ZINB-WaVE | 256.19 | 40 | 530 |

**Table 3:** Time and RAM usage of NewWave on the 1.3 million cell dataset. This table reports the data displayed in Fig. 1D.

| **Model** | **# of cells** | **RAM in GB** | **Time in minutes** |
| --- | --- | --- | --- |
| hdf5 | 10000 | 9 | 28.364 |
| hdf5_40cores | 10000 | 33 | 9.063 |
| matrix | 10000 | 5 | 3.768 |
| pca | 10000 | 1 | 5.452 |
| hdf5 | 100000 | 12 | 98.240 |
| hdf5_40cores | 100000 | 40 | 31.081 |
| matrix | 100000 | 26 | 36.611 |
| pca | 100000 | 5 | 13.375 |
| hdf5 | 200000 | 16 | 153.370 |
| hdf5_40cores | 200000 | 46 | 49.374 |
| matrix | 200000 | 61 | 79.583 |
| pca | 200000 | 9 | 19.562 |
| hdf5 | 300000 | 19 | 222.784 |
| hdf5_40cores | 300000 | 48 | 68.381 |
| matrix | 300000 | 98 | 132.788 |
| pca | 300000 | 11 | 30.691 |
| hdf5 | 600000 | 25 | 372.509 |
| hdf5_40cores | 600000 | 55 | 124.352 |
| matrix | 600000 | 197 | 198.116 |
| pca | 600000 | 19 | 51.960 |
| hdf5 | 1300000 | 55 | 730.297 |
| hdf5_40cores | 1300000 | 109 | 271.085 |
| matrix | 1300000 | 419 | 494.515 |
| pca | 1300000 | 40 | 102.940 |

## References

Pagès, H., with contributions from Peter Hickey, and Lun, A. (2019). DelayedArray: Delayed operations on array-like objects.

Raimundo, F., Vallot, C., and Vert, J.-P. (2020). Tuning parameters of dimensionality reduction methods for single-cell rna-seq analysis. Genome biology, 21(1), 1–17.

Risso, D., Perraudeau, F., Gribkova, S., Dudoit, S., and Vert, J. P. (2018). A general and flexible method for signal extraction from single-cell RNA-seq data. Nature Communications.

Sun, S., Zhu, J., Ma, Y., and Zhou, X. (2019). Accuracy, robustness and scalability of dimensionality reduction methods for single-cell rna-seq analysis. Genome Biology, 20(1), 1–21.

Svensson, V. (2020). Droplet scrna-seq is not zero-inflated. Nature Biotechnology, 38(2), 147–150.

Townes, F. W., Hicks, S. C., Aryee, M. J., and Irizarry, R. A. (2019). Feature selection and dimension reduction for single-cell rna-seq based on a multinomial model. Genome Biology, 20(1), 295.

Yao, Z., Liu, H., Xie, F., Fischer, S., Sina Booeshaghi, A., Adkins, R. S., Aldridge, A. I., Ament, S. A., Pinto-Duarte, A., Bartlett, A., Margarita Behrens, M., van den Berge, K., Bertagnolli, D., Biancalani, T., Corrada Bravo, H., Casper, T., Colantuoni, C., Creasy, H., Crichton, K., Crow, M., Dee, N., Dougherty, E. L., Doyle, W. I., Dudoit, S., Fang, R., Felix, V., Fong, O., Giglio, M., Goldy, J., Hawrylycz, M., Roux de Bézieux, H., Herb, B. R., Hertzano, R., Hou, X., Hu, Q., Crabtree, J., Kancherla, J., Kroll, M., Lathia, K., Li, Y. E., Lucero, J. D., Luo, C., Mahurkar, A., McMillen, D., Nadaf, N., Nery, J. R., Niu, S. Y., Orvis, J., Osteen, J. K., Pham, T., Poirion, O., Preissl, S., Purdom, E., Rimorin, C., Risso, D., Rivkin, A. C., Smith, K., Street, K., Sulc, J., Nguyen, T. N., Tieu, M., Torkelson, A., Tung, H., Vaishnav, E. D., Svensson, V., Vanderburg, C. R., Ntranos, V., van Velthoven, C., Wang, X., White, O. R., Josh Huang, Z., Kharchenko, P. V., Pachter, L., Ngai, J., Regev, A., Tasic, B., Welch, J. D., Gillis, J., Macosko, E. Z., Ren, B., Ecker, J. R., Zeng, H., and Mukamel, E. A. (2020). An integrated transcriptomic and epigenomic atlas of mouse primary motor cortex cell types. biorXiv.

## References

Davide Risso, Fanny Perraudeau, Svetlana Gribkova, Sandrine Dudoit, and Jean Philippe Vert. A general and flexible method for signal extraction from single-cell RNA-seq data. Nature Communications, 2018. ISSN 20411723. doi: 10.1038/s41467-017-02554-5.

Zizhen Yao, Hanqing Liu, Fangming Xie, Stephan Fischer, A. Sina Booeshaghi, Ricky S Adkins, Andrew I Aldridge, Seth A Ament, Antonio Pinto-Duarte, Anna Bartlett, M. Margarita Behrens, Koen van den Berge, Darren Bertagnolli, Tommaso Biancalani, Héctor Corrada Bravo, Tamara Casper, Carlo Colantuoni, Heather Creasy, Kirsten Crichton, Megan Crow, Nick Dee, Elizabeth L Dougherty, Wayne I Doyle, Sandrine Dudoit, Rongxin Fang, Victor Felix, Olivia Fong, Michelle Giglio, Jeff Goldy, Mike Hawrylycz, Hector Roux de Bézieux, Brian R Herb, Ronna Hertzano, Xiaomeng Hou, Qiwen Hu, Jonathan Crabtree, Jayaram Kancherla, Matthew Kroll, Kanan Lathia, Yang Eric Li, Jacinta D Lucero, Chongyuan Luo, Anup Mahurkar, Delissa McMillen, Naeem Nadaf, Joseph R Nery, Sheng Yong Niu, Joshua Orvis, Julia K Osteen, Thanh Pham, Olivier Poirion, Sebastian Preissl, Elizabeth Purdom, Christine Rimorin, Davide Risso, Angeline C Rivkin, Kimberly Smith, Kelly Street, Josef Sulc, Thuc Nghi Nguyen, Michael Tieu, Amy Torkelson, Herman Tung, Eeshit Dhaval Vaishnav, Valentine Svensson, Charles R Vanderburg, Vasilis Ntranos, Cindy van Velthoven, Xinxin Wang, Owen R White, Z. Josh Huang, Peter V Kharchenko, Lior Pachter, John Ngai, Aviv Regev, Bosiljka Tasic, Joshua D Welch, Jesse Gillis, Evan Z Macosko, Bing Ren, Joseph R Ecker, Hongkui Zeng, and Eran A Mukamel. An integrated transcriptomic and epigenomic atlas of mouse primary motor cortex cell types. biorXiv, 2020. URL https://doi.org/10.1101/2020.02.29.970558.

Laleh Haghverdi, Aaron TL Lun, Michael D Morgan, and John C Marioni. Batch effects in single-cell rna-sequencing data are corrected by matching mutual nearest neighbors. Nature biotechnology, 36(5):421–427, 2018.

